# Lyophilized *B. subtilis* ZB183 Spores: 90-Day Repeat Dose Oral (Gavage) Toxicity Study In Wistar Rats

**DOI:** 10.1101/724542

**Authors:** Appala Naidu. B, Kamala Kannan, D. P. Santhosh Kumar, John W.K. Oliver, Zachary D. Abbott

**Affiliations:** Department of Safety Assessment, Eurofins Advinus Limited, Post Box No. 5813, Plot Nos. 21 & 22 Peenya Ii Phase, Bengaluru 560 058, India; ZBiotics Company, 181 2nd St., San Francisco, CA 94105 USA

## Abstract

A 90-day repeated-dose oral toxicological evaluation was conducted according to GLP and OECD guidelines on lyophilized spores of the novel genetically modified strain *B. subtilis* ZB183. Lyophilized spores at doses of 10^9^, 10^10^, and 10^11^ CFU/kg body weight/day were administered by oral gavage to Wistar rats for a period of 90 consecutive days. *B.Subtilis* ZB183 had no effects on clinical signs, mortality, ophthalmological examinations, functional observational battery, body weights, body weight gains and food consumption in both sexes. There were no test item-related changes observed in haematology, coagulation, urinalysis, thyroid hormonal analysis, terminal fasting body weights, organ weights, gross pathology and histopathology. A minimal increase in the plasma albumin level was observed at 10^10^ and 10^11^ CFU/kg/day doses without an increase in total protein in males or females and was considered a non-adverse effect. The “No Observed Adverse Effect Level (NOAEL)” is defined at the highest dose of 10^11^ CFU/kg body weight/day for lyophilized *B.Subtilis* ZB183 Spores under the test conditions employed.

## Introduction

*B. subtilis* is a gram-positive, rod-shaped, endospore-forming bacterium found in the soil, water sources, associated with plants, and in the gastrointestinal tract of humans [1,2]. *B. subtilis* strains have a long history of human consumption. For example, *B. subtilis* is used in the traditional Japanese fermented soybean dish called natto, with a bacterial concentration reported as approximately 10^8^ CFU/g [1,3]. *B. subtilis* strains have also been sold in numerous probiotic products around the globe, at levels of 10^6^–10^9^ CFU/serving [3]. *B. subtilis* is also listed in the International Dairy Federation’s (originally a collaboration with the European Food and Feed Culture’s Association) inventory of microbial species with technological beneficial roles in fermented food products, emphasizing its long history of use [4].

ZB183 is a viable genetically modified *B. subtilis* strain that constitutively expresses an acetaldehyde dehydrogenase enzyme, AcoD, from *C. necator* (which has been known by a number of other names, including *Alcaligenes eutrophus*, *Ralstonia eutropha*, and *Hydrogenomonas eutropha*). The parental strain is *B. subtilis* PY79, a widely used strain in laboratory studies with a published sequence, that is lacking of many mobile genetic elements found in other *B. subtilis* strains, and has been found to be nontoxic to vertebrates and has been studied in humans [5–7].

The mechanism of action of the acetaldehyde dehydrogenase enzyme is to oxidize acetaldehyde to acetic acid using the cofactor NAD+. Aldehyde dehydrogenase enzymes are ubiquitously found in all three taxonomic domains (Archaea, Eubacteria and Eukarya), and a common ancestral gene dating back ∼ 3 billion years for these enzymes has been suggested [8–11]. Members of this “superfamily” of enzymes aid in the prevention of toxic accumulation of aldehydes from endogenous production (e.g. metabolism of amino acids, carbohydrates, lipids, and more) or from exogenous exposures (e.g. alcohol consumption and metabolism) [8,9,12,13].

Aldehydes are found in various food substances, for example as products of food processing or as flavoring additives (e.g. anisaldehyde, vanillin, and citral), in various milk products, fruits/vegetables, and as a breakdown product of alcohol consumption, and they may exhibit toxicity due to their chemical reactivity [10,14–16]. Aldehyde dehydrogenase enzymes are found in the human epithelium of the gastrointestinal tract (e.g. in the saliva and stomach), and certain food substances such as sulforaphane in cruciferous vegetables have been shown to induce these endogenous enzymes [17,18]. Detoxification of aldehydes found in foods and beverages by bacteria expressing aldehyde dehydrogenase enzymes, which function under desirable conditions such as certain pH levels, has been the subject of investigations[10,19].

The donor species of the AcoD acetaldehyde dehydrogenase enzyme, *C. necator* [20–22] is a gram-negative betaproteobacterium, which has been well studied for its ability to store large amounts of organic carbon that can be used as biodegradable plastic [23]. It has also been studied for its nutritive value, due to its high protein content and quality (e.g. for animal feed) [24]. The sequence of the AcoD is published and the amino acid sequence is similar to other acetaldehyde dehydrogenases; it is 44.5% identical to that found in human liver, 23.6% to *Saccharomyces cerevisiae*, 41.1% to rat liver, and 40.0% to *Escherichia coli* [20].

The safety of the parent strain *B. subtilis* PY79 is well established. Acetaldehyde dehydrogenases are ubiquitous in nature and their safety is also well established. Additionally, the ingestion of enzymes is generally considered to be safe and unlikely to be allergenic [25]. However, the combination of the two in the novel strain *B. subtilis* ZB183 has not been evaluated. The objective of this study was to assess the toxicological profile and an estimate of the No Observed Effect Level (NOEL) / No Observed Adverse Effect Level (NOAEL) of lyophilized *B.Subtilis* ZB183 Spores, when administered by oral gavage to Wistar rats for a period of 90 consecutive days.

## Methods

The study was performed in compliance with the OECD Principles of Good Laboratory Practice [C (97) 186/Final] and US FDA Good Laboratory Practice for Nonclinical Laboratory Studies (21 CFR Part 58). This study was performed in an Association for Assessment and Accreditation of Laboratory Animal Care International (AAALAC) (http://www.aaalac.org), accredited facility. All procedures were in compliance with the guidelines provided by the Committee for Purpose of Control and Supervision of Experiments on Animals (CPCSEA) India. This study plan has been approved by the Institutional Animal Ethics Committee (IAEC) of Eurofins Advinus Limited. The study of *B. Subtilis* ZB183 spores was conducted as per OECD Guideline No. 408 for Testing of Chemicals, “Repeated Dose 90 Day Oral Toxicity Study in Rodents” adopted on 25 June, 2018.

Two lots of lyophilized *B. Subtilis* ZB183 spores were provided by the manufacturer, The Saskatchewan Research Council, 125 – 15 Innovation Blvd. Saskatoon, SK S7N 2X8 Canada. Lyophilized spores were provided as a clumpy dark brown powder. Two lots of test articles (300L engineering batch ZBT-002a and 300L engineering batch ZBT-002a part 2) were used in the study and both batches were made according to commercial GMP specifications and spores were assayed for genetic strain purity by PCR and standard microbial enumeration techniques to identify any contaminating coliforms, yeast or mold. Heavy metal content assayed by ICP-MS was below baseline. Samples were retested at the end of the study to confirm the purity and viability of the spores was maintained throughout the testing period.

Based on prior testing of probiotic *B. subtilis*, no safety concerns were expected even at high doses [26–32]. Therefore, dosage levels for the study were chosen based on the intended dose for the final product, namely 10^9^ CFU. 10- and 100-fold higher dosage levels were tested to determine if the NOAEL was at least 100-fold higher than the intended dose. The target doses were calculated and administered based on the colony forming units (CFU) count per weight of the lyophilized spore material. The CFU count of each batch was measured by standard serial dilution and plating on LB agar plates. In brief, spore suspension is serially diluted and plated on rich media agar. The number of colonies formed was counted and adjusted by the dilution factor to calculate the initial concentration.

The test article doses were prepared as per Table 1 by suspending lyophilized spores in MilliQ water to achieve concentrations of 2, 20, and 200mg/mL for the first half of the lot and 1, 10, and 100mg/mL for the second half of the lot with a consistent dose volume of 10 mL/kg body weight. The second half of the lot spent more time in the lyophilizer and thus had lower moisture content. Thus, the concentration of CFU/g was higher and less material was needed to get comparable doses of CFU. Both halves of the lot were individually tested for purity and stability as discussed above. The first half of the lot was used to dose the animals from days 1-66, and the second half of the lot was used from days 67-90. The dose formulations were prepared daily prior to administration of the test item and the control group received the same volume of 100% MilliQ Water. Dose formulations were analyzed for concentration of viable bacterial cells on Day 1, 44, 67, and 84.

**TABLE 1.**
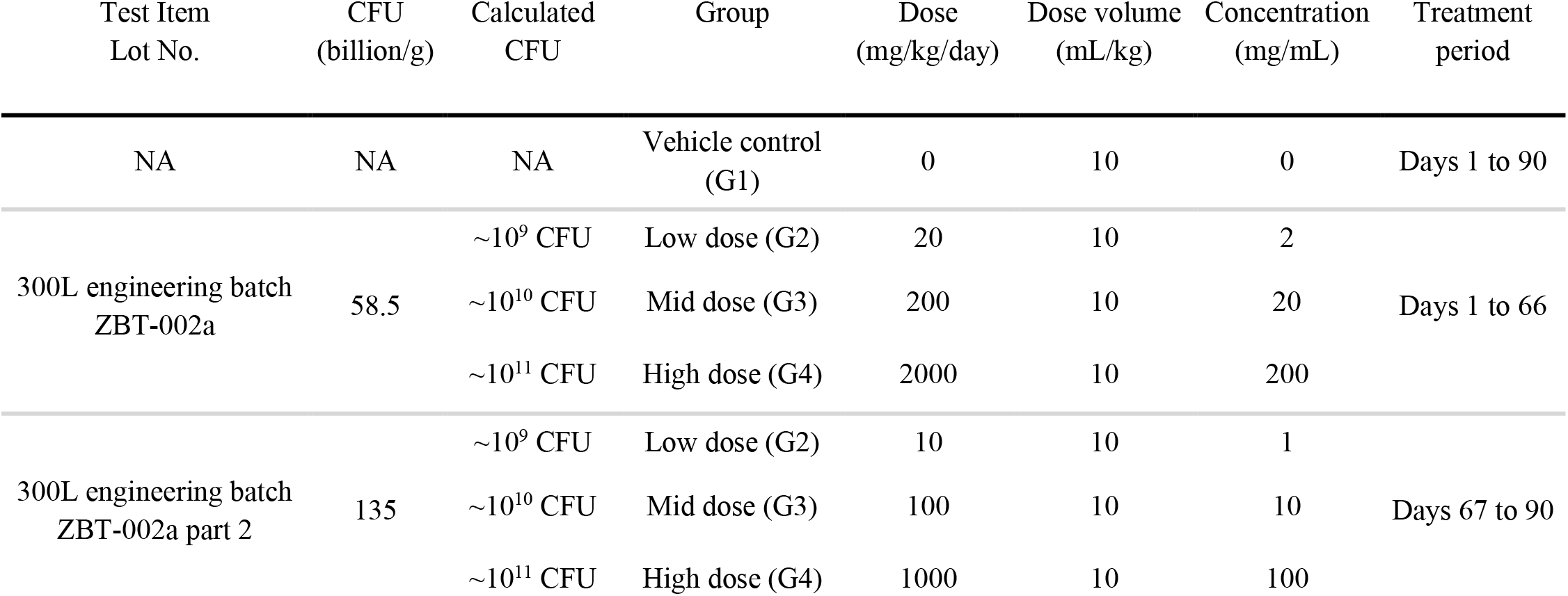
Details of CFU/g for both lot numbers of test item, doses administered and treatment days.

The vehicle control and test item treated groups each consisted of 10 rats/sex. Two rats per cage/sex/group were housed in standard polysulfone cages (size: Length 425 × Breadth 266 × Height 185 mm), with stainless steel top grill having facilities for pelleted food and polycarbonate drinking water bottle with stainless steel sipper tubes. Steam sterilized corn cob was used as bedding and changed along with the cage at least once a week. Teklad Certified (2014C) Global 14% Protein Rodent Maintenance Diet – pellet (Certified) manufactured by Envigo PO Box 44220, Madison, WI 53744-4220, was provided *ad libitum* to rats. Deep bore-well water passed through activated charcoal filter and exposed to UV rays in ‘Aquaguard’ an on-line water filter-purifier manufactured by Eureka Forbes Ltd., Mumbai 400 001, India, was provided *ad libitum* to rats in polycarbonate bottles with stainless steel sipper tubes. Rats were housed under standard laboratory conditions, air conditioned with adequate fresh air supply (12 - 15 air changes/hour), temperature in the range of 20 to 24 °C, relative humidity between 58 and 67 %, and with 12 hours light and 12 hours dark cycle.

Rats were randomly distributed to different groups by the body weight stratification method using Provantis™ Software (Version 10.1.0.1, Instem LSS, Staffordshire ST15OSD, United Kingdom) prior to the start of treatment. Dose formulations and vehicle were administered by oral gavage to the specific treatment and vehicle control groups, respectively, once daily at approximately the same time each day (varied by ± 3 hours) for 90 consecutive days. The dose volume administered was at an equivolume of 10 mL/kg and was calculated for individual animal on the first day of the treatment and was adjusted according to the recent body weights recorded at different intervals of the study.

In this study, assessments included: clinical signs, detailed clinical observations, ophthalmological observations, neurological observations, mortality, body weight, food consumption, functional observation battery, haematology, clinical chemistry, urinalysis, gross pathology, organ weights and histopathological evaluation. All rats were observed for clinical signs twice a day (pre-dose and post-dose) during treatment days. The observation for morbidity and mortality was carried out twice daily. As there were no clinical signs of concern, the observation for morbidity and mortality was carried out once during weekends and public holidays. Detailed clinical examination was carried out prior to the test item administration on Day 1 and at weekly intervals thereafter (±1 day) during the treatment period.

Ophthalmological examination of all animals was carried out prior to start of treatment and at the end of the treatment for all the groups (Day 90). Before examination, mydriasis was induced using a 1 % solution of Tropicamide.

Standard neurological examination was conducted during week 13 of the treatment period for all rats in the toxicity groups to assess behavioral and neurological status of rats. The objective of neurological examination was to observe the subject’s response to handling and conducted other procedures of the neurological examinations that could best performed when the rat was being held. Each rat was observed for the following examinations: Ease of removal from home cage, handling reactivity, palpebral closure, eye examination, piloerection, lacrimation, salivation, skin/fur examination, perineum wetness, respiration, muscle tone, and extensor thrust response. The observations were recorded using scores. In addition, each rat was placed in an open field arena and observed for the following: gait, posture, tremors, mobility score, arousal level, clonic or tonic movements, stereotypic behaviour, bizarre behaviour, urination, defecation, abnormal vocalizations, and rearing. Again, the observations were recorded using scores.

Each rat was also tested in a functional observation battery, including sensory evaluation, measurement of grip performance, landing hindlimb foot splay, rectal temperature and motor activity. For the Landing Hindlimbs Footsplay test, the hind feet of the animal was marked with ink. The rat was suspended in a prone position and then dropped from a height of approximately 30 cm on to a recording format. A total of 3 trials of the distance (in mm) between the centre of the back of the heel prints were recorded for each rat. A clean recording paper was used for each rat and 3 footsplay readings were presented in the report.

The motor activity of rats was measured using an automated animal activity measuring system (Columbus Instruments, USA) equipped with a computer analyzer. Each rat was individually placed in the activity cages of the instrument. The rats were monitored for 30 minutes. During this motor activity measurement session, parameters viz., the stereotypic time (small movements) in seconds, the ambulatory time (large ambulatory movement) in seconds, horizontal counts and ambulatory counts were monitored. The Opto-Varimex 4 motor activity measurement system provides the data at 1 minute intervals and the data was analyzed in blocks of 10 minutes intervals and presented in the report. Body temperature (rectal temperature) was measured at the time of functional test.

Individual body weights were recorded prior to test item administration on Day 1 and weekly thereafter (± 1 day) for all groups of rats during the treatment period. Fasting body weight was recorded prior to sacrifice on Day 91. Blood samples (approximately 3 mL) were collected at the end of the treatment period (Day 91) from all rats under isoflurane anesthesia, with a fine capillary tube, by retro-orbital sinus puncture. Aliquots of blood for clinical pathology tests were collected into tubes containing anticoagulants as follows: Haematology 0.7 ml blood with K_2_EDTA (1.6 mg/ml), Clinical chemistry 1.8 ml blood with Lithium heparin (10 Units/ml) and Coagulation 0.5 ml blood with Trisodium citrate (3.2 mg/ml).

The following haematological parameters were determined using ADVIA 2120i haematology system (Siemens Healthcare Diagnostics Inc., NY, USA): red blood corpuscles, RBC, 10^12^/L; haemoglobin, HGB, g/L; haematocrit, HCT, L/L; mean corpuscular volume, MCV, fL; mean corpuscular haemoglobin, MCH, pg; mean corpuscular haemoglobin concentration, MCHC, g/L; reticulocytes count, Retic, 10^12^/L & %; white blood corpuscles, WBC, 10^9^/L; differential leukocyte count (differential leukocyte parameters and their respective abbreviations are: neutrophils [Neut], lymphocytes [Lymp], monocytes [Mono], eosinophils [Eosi] and basophils [Baso]), DLC, 10^9^/L; platelets (Plat), 10^9^/L; red blood cell morphology (red cell distribution width, haemoglobin distribution width, hyperchromic cells, hypochromic cells, macrocytes, microcytes, RBC fragments, RBC ghosts).

For thyroid hormone analysis, blood samples were collected from all rats along with blood collection for clinical pathology investigation prior to sacrifice for the determination of total T4, T3 and TSH hormones in serum by enzyme-linked immunosorbent assay (ELISA). Urine samples were collected on Day 91 from all rats.

All rats in the study were subjected to detailed necropsy on Day 91 (examination of external surfaces of the body, all orifices; cranial, thoracic and abdominal cavities and their contents). On completion of the gross pathology examination, the tissues and organs (adrenal glands, aorta, bone marrow smear, brain (cerebrum, cerebellum, medulla/pons), cecum, cervix, colon, duodenum, epididymides, esophagus, eyes, femur bone with distal joint, gross lesions, gut associated lymphoid tissue, glands, harderian, heart, ileum, jejunum, kidneys, larynx, liver, lungs, axillary lymph node, mesenteric lymph node, mandibular lymph node, mammary gland, optic nerve, sciatic nerve, nose, ovaries, oviducts, pancreas, pharynx, pituitary, prostate, rectum, salivary glands, seminal vesicles with coagulating glands, skeletal muscle, skin, spinal cord, spleen, sternum with marrow, stomach, testes, thymus, thyroid with parathyroid, tongue, trachea, ureters, urinary bladder, uterus, and vagina.) were collected and weighed from all rats. The organ weight ratios as percentage of fasting body weight and brain weight were determined and presented in the report. The paired organs were weighed together and combined weights were presented. The tissues were preserved in 10% Neutral Buffered Formalin (NBF) except for the testes (modified Davidson fluid) and eyes (Davidson fluid). A full histopathological examination was carried out on the preserved organs of the vehicle control (G1) and high dose (G4) group rats and on all gross lesions. Tissues from lower dose groups were not examined as there were no test item-related microscopic changes at high dose. The tissues were processed for routine paraffin embedding and approximately 5 micron sections were stained with Mayer’s haematoxylin and eosin stain.

Statistical analysis was performed on data captured using Provantis™: Parameters such as body weight, organ weights, laboratory investigations - haematology (coagulation parameters data were directly captured in Provantis™) and clinical chemistry data were analysed using Provantis™ built-in statistical tests. Derived data like net body weight gains, food consumption and organ weight ratios was also analysed using Provantis™ built-in statistical tests. For data captured outside of Provantis™, the statistical analysis of the experimental data was carried out using licensed copies of SYSTAT Statistical package Ver.12.0. The T3, T4, TSH data were tested for normality (Shapiro-Wilk test) and homogeneity of variances (Levene’s test) within the group before performing a one-factor ANOVA. All quantitative variables of neurological observations (neuromuscular observation/body temperature/body weights) were tested for normality (Shapiro-Wilk test) and homogeneity of variances (Levene’s test) within the group before performing a one-factor ANOVA modeling by treatment groups. Non-optimal (non-normal or heteroscedastic) data was transformed, before ANOVA was performed. Comparison of means between treatment groups and the vehicle control group was done using Dunnett’s ‘t’ test when the overall treatment ‘F’ test was found significant. All analyses and comparisons were evaluated at the 5% (p < 0.05) level.

## Results and Discussion

There were no mortalities at any of the tested doses throughout the treatment period. There were no clinical signs in the vehicle control group and in all the test item treated groups throughout the treatment period. Ophthalmological examination at the start and end of treatment for all rats did not reveal any abnormalities in the eyes. There were no test item-related neurological abnormalities observed in home cage, while handling, during open field, or sensory observations.

An incidence of statistically significant lower hindlimb foot splay was observed in mid-dose males (Table 2). This result does not follow a dose-relationship and is considered as incidental. There were no test item-related variations observed in forelimbs and hind limbs grip strength. There were no test item-related variations observed in the motor activity.

**TABLE 2.**
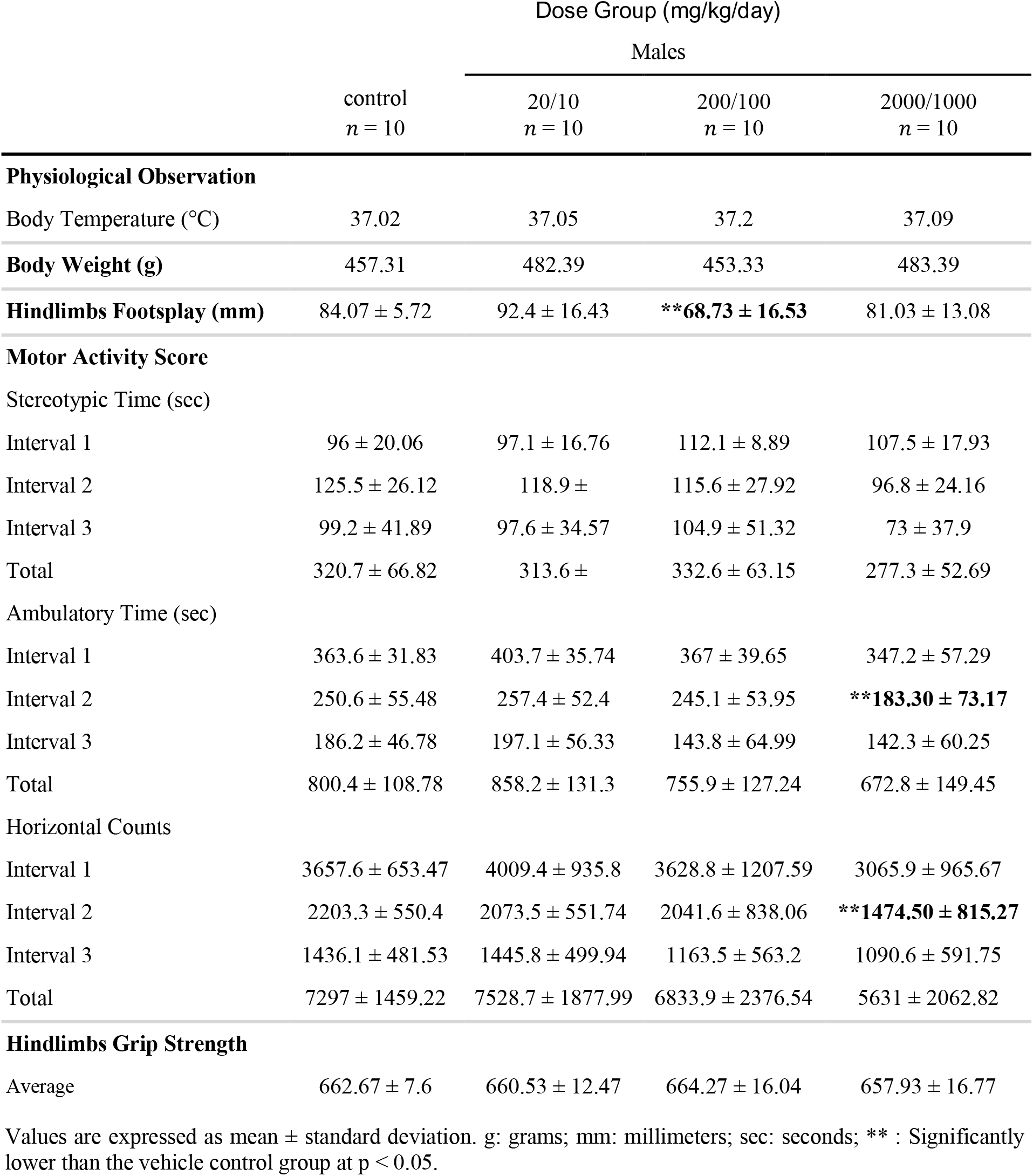
Summary of Significant Functional Observation Battery Results.

|  |  | Dose Group (mg/kg/day) |  |  |
| --- | --- | --- | --- | --- |
|  |  | Males |  |  |
|  | control<br>*n* = 10 | 20/10<br>*n* = 10 | 200/100<br>*n* = 10 | 2000/1000<br>*n* = 10 |
| **Physiological Observation** |  |  |  |  |
| Body Temperature (°C) | 37.02 | 37.05 | 37.2 | 37.09 |
| **Body Weight (g)** | 457.31 | 482.39 | 453.33 | 483.39 |
| **Hindlimbs Footsplay (mm)** | 84.07 ± 5.72 | 92.4 ± 16.43 | **\*\*68.73 ± 16.53** | 81.03 ± 13.08 |
| **Motor Activity Score** |  |  |  |  |
| Stereotypic Time (sec) |  |  |  |  |
| Interval 1 | 96 ± 20.06 | 97.1 ± 16.76 | 112.1 ± 8.89 | 107.5 ± 17.93 |
| Interval 2 | 125.5 ± 26.12 | 118.9 ± | 115.6 ± 27.92 | 96.8 ± 24.16 |
| Interval 3 | 99.2 ± 41.89 | 97.6 ± 34.57 | 104.9 ± 51.32 | 73 ± 37.9 |
| Total | 320.7 ± 66.82 | 313.6 ± | 332.6 ± 63.15 | 277.3 ± 52.69 |
| Ambulatory Time (sec) |  |  |  |  |
| Interval 1 | 363.6 ± 31.83 | 403.7 ± 35.74 | 367 ± 39.65 | 347.2 ± 57.29 |
| Interval 2 | 250.6 ± 55.48 | 257.4 ± 52.4 | 245.1 ± 53.95 | **\*\*183.30 ± 73.17** |
| Interval 3 | 186.2 ± 46.78 | 197.1 ± 56.33 | 143.8 ± 64.99 | 142.3 ± 60.25 |
| Total | 800.4 ± 108.78 | 858.2 ± 131.3 | 755.9 ± 127.24 | 672.8 ± 149.45 |
| Horizontal Counts |  |  |  |  |
| Interval 1 | 3657.6 ± 653.47 | 4009.4 ± 935.8 | 3628.8 ± 1207.59 | 3065.9 ± 965.67 |
| Interval 2 | 2203.3 ± 550.4 | 2073.5 ± 551.74 | 2041.6 ± 838.06 | **\*\*1474.50 ± 815.27** |
| Interval 3 | 1436.1 ± 481.53 | 1445.8 ± 499.94 | 1163.5 ± 563.2 | 1090.6 ± 591.75 |
| Total | 7297 ± 1459.22 | 7528.7 ± 1877.99 | 6833.9 ± 2376.54 | 5631 ± 2062.82 |
| **Hindlimbs Grip Strength** |  |  |  |  |
| Average | 662.67 ± 7.6 | 660.53 ± 12.47 | 664.27 ± 16.04 | 657.93 ± 16.77 |

Values are expressed as mean ± standard deviation. g: grams; mm: millimeters; sec: seconds; \*\*: Significantly lower than the vehicle control group at $p < 0.05$ .

A small number of statistically significant changes observed in motor activity parameters were randomly observed at different intervals across the treatment groups in males as compared to the vehicle control group (Table 2). These changes were inconsistent and did not show any dose-dependent pattern. There were no test item-related variations observed in the body temperature (°C) in all the test item treated groups.

There were no test item-related variations observed in the body weights (Table 2). Additionally, there were no test item-related variations in food consumption in all tested dose groups in both sexes during the treatment period. There were no test item-related changes in hematology parameters. An increase in basophil count in females (Table 3) and decrease in hyperchromic cells in males (Table 3) at the high dose were observed but were not accompanied by changes in other leukocyte counts and RBC parameters and thus considered as incidental. The prothrombin time and APTT values were not significantly affected by test item administration at all the tested doses.

**TABLE 3.** Summary of Significant Hematology and Coagulation parameters.

|  | Dose Group (mg/kg/day) |  |  |  |  |  |  |  |
| --- | --- | --- | --- | --- | --- | --- | --- | --- |
|  | Males |  |  |  | Females |  |  |  |
|  | control<br>*n* = 10 | 20/10<br>*n* = 10 | 200/100<br>*n* = 10 | 2000/1000<br>*n* = 10 | control<br>*n* = 10 | 20/10<br>*n* = 10 | 200/100<br>*n* = 10 | 2000/1000<br>*n* = 10 |
| RBC ($10^{12}/L$) | 8.94 ± 0.36 | 8.64 ± 0.29 | 9.00 ± 0.47 | 8.83 ± 0.34 | 8.21 ± 0.24 | 8.41 ± 0.34 | 8.10 ± 0.28 | 8.23 ± 0.31 |
| HGB (g/L) | 150 ± 5 | 146 ± 2 | 150 ± 6 | 149 ± 4 | 149 ± 5 | 149 ± 5 | 148 ± 4 | 149 ± 5 |
| HCT (L/L) | 0.467 ±<br>0.015 | 0.454 ±<br>0.015 | 0.468 ±<br>0.021 | 0.464 ±<br>0.012 | 0.458 ±<br>0.012 | 0.461 ±<br>0.015 | 0.456 ±<br>0.016 | 0.459 ±<br>0.013 |
| MCV (fL) | 52.2 ± 1.2 | 52.6 ± 1.7 | 52.0 ± 1.8 | 52.6 ± 1.3 | 55.8 ± 1.1 | 54.0 ± 1.5 | 56.2 ± 2.0 | 55.8 ± 1.3 |
| MCH (pg) | 16.8 ± 0.6 | 17.0 ± 0.7 | 16.7 ± 0.6 | 16.0 ± 0.6 | 18.2 ± 0.5 | 17.7 ± 0.6 | 18.3 ± 0.7 | 18.1 ± 0.5 |
| MCHC (g/L) | 323 ± 6 | 323 ± 9 | 321 ± 4 | 321 ± 7 | 325 ± 3 | 323 ± 6 | 325 ± 7 | 325 ± 5 |
| Retic A ($10^{12}/L$) | 0.183 ±<br>0.035 | 0.201 ±<br>0.022 | 0.205 ±<br>0.022 | 0.193 ±<br>0.032 | 0.198 ±<br>0.030 | 0.196 ±<br>0.034 | 0.200 ±<br>0.027 | 0.180 ±<br>0.025 |
| Retic (%) | 2.05 ± 0.37 | 2.32 ± 0.22 | 2.28 ± 0.23 | 2.20 ± 0.40 | 2.42 ± 0.41 | 2.34 ± 0.42 | 2.47 ± 0.34 | 2.18 ± 0.28 |
| RDW (%) | 13.2 ± 0.4 | 14.0 ± 1.4 | 13.5 ± 0.6 | 13.4 ± 0.4 | 11.7 ± 0.4 | 11.9 ± 0.4 | 11.9 ± 0.5 | 11.6 ± 0.4 |
| HDW (g/L) | 25.2 ± 1.0 | 27.1 ± 4.2 | 26.5 ± 1.2 | 24.6 ± 0.9 | 21.1 ± 1.0 | 21.3 ± 0.6 | 21.4 ± 0.4 | 20.6 ± 0.6 |
| Hyper (%) | 2.9 ± 0.7 | 3.2 ± 2.0 | 2.9 ± 0.6 | **\*\*1.7 ± 0.4** | 1.2 ± 0.3 | 1.1 ± 0.5 | 1.2 ± 0.7 | 1.0 ± 0.3 |
| Hypo (%) | 0.1 ± 0.0 | 0.1 ± 0.0 | 0.1 ± 0.0 | 0.1 ± 0.0 | 0.1 ± 0.0 | 0.1 ± 0.0 | 0.1 ± 0.0 | 0.1 ± 0.0 |
| Macro (%) | 0.0 ± 0.0 | 0.1 ± 0.2 | 0.0 ± 0.0 | 0.0 ± 0.0 | 0.0 ± 0.0 | 0.0 ± 0.0 | 0.0 ± 0.0 | 0.0 ± 0.0 |
| Micro (%) | 0.0 ± 0.0 | 0.0 ± 0.0 | 0.0 ± 0.0 | 0.0 ± 0.0 | 0.0 ± 0.0 | 0.0 ± 0.0 | 0.0 ± 0.0 | 0.0 ± 0.0 |
| RBC Fragments<br>($10^{12}/L$) | 0.08 ± 0.01 | 0.09 ± 0.01 | 0.09 ± 0.01 | 0.09 ± 0.01 | 0.07 ± 0.02 | 0.08 ± 0.01 | 0.07 ± 0.01 | 0.08 ± 0.02 |
| RBC Ghosts<br>($10^{12}/L$) | 0.04 ± 0.02 | 0.03 ± 0.01 | 0.03 ± 0.01 | 0.03 ± 0.01 | 0.03 ± 0.00 | 0.03 ± 0.01 | 0.04 ± 0.01 | 0.03 ± 0.01 |
| PLT ($10^9/L$) | 962 ± 88 | 972 ± 74 | 1010 ± 138 | 1037 ± 61 | 938 ± 135 | 984 ± 90 | 945 ± 62 | 1027 ± 101 |
| WBC ($10^9/L$) | 6.50 ± 0.89 | 6.59 ± 1.43 | 6.42 ± 1.28 | 5.53 ± 1.85 | 3.58 ± 1.12 | 3.77 ± 0.79 | 3.50 ± 0.62 | 4.34 ± 1.14 |
| Neut A ($10^9/L$) | 1.15 ± 0.36 | 1.15 ± 0.24 | 1.40 ± 0.44 | 0.93 ± 0.33 | 0.76 ± 0.23 | 0.40 ± 0.37 | 0.75 ± 0.26 | 0.72 ± 0.16 |
| Lymp A ($10^9/L$) | 5.05 ± 0.68 | 5.10 ± 1.59 | 4.61 ± 1.04 | 4.34 ± 1.49 | 2.59 ± 0.98 | 2.50 ± 0.58 | 2.56 ± 0.54 | 3.43 ± 1.02 |
| Mono A ($10^9/L$) | 0.22 ± 0.07 | 0.24 ± 0.07 | 0.29 ± 0.13 | 0.18 ± 0.07 | 0.13 ± 0.03 | 0.15 ± 0.05 | 0.11 ± 0.05 | 0.10 ± 0.04 |
| Baso A ($10^9/L$) | 0.01 ± 0.00 | 0.01 ± 0.01 | 0.01 ± 0.01 | 0.01 ± 0.01 | 0.00 ± 0.00 | 0.00 ± 0.00 | 0.01 ± 0.01 | **\*0.01 ± 0.00** |
| Eosi A (10<sup>9</sup>/L) | 0.07 ± 0.03 | 0.07 ± 0.03 | 0.10 ± 0.04 | 0.08 ± 0.04 | 0.09 ± 0.05 | 0.11 ± 0.09 | 0.07 ± 0.04 | 0.08 ± 0.03 |
| Prothrombin Time (seconds) | 17.1 ± 0.2 | 16.5 ± 0.4 | **\*\*15.7 ± 0.6** | 17.0 ± 0.9 | 16.1 ± 0.7 | 16.1 ± 0.7 | 15.8 ± 0.5 | **\*\*15.4 ± 0.5** |
| APTT (seconds) | 15.8 ± 1.6 | **\*\*11.8 ± 2.3** | **\*\*13.4 ± 1.7** | 13.8 ± 1.8 | 15.2 ± 2.5 | 13.1 ± 1.5 | 15.1 ± 1.8 | 14.1 ± 1.8 |

Values are expressed as mean ± standard deviation. mg/kg, milligrams per kilogram body weight per day; L, liter; n, number of animals; \*/\*\* Significantly higher/lower than the vehicle control group at p < 0.05

A minimal increase in plasma albumin was present at the mid and high doses and considered as test item-related. However, the corresponding total protein levels were not changed significantly at the high dose (Table 4). In the absence of elevated protein, increased albumin is considered a non-adverse finding.

**TABLE 4.** Summary of Clinical Chemistry Parameters.

|  | Dose Group (mg/kg/day) |  |  |  |  |  |  |  |
| --- | --- | --- | --- | --- | --- | --- | --- | --- |
|  | Males |  |  |  | Females |  |  |  |
|  | control<br>n = 10 | 20/10<br>n = 10 | 200/100<br>n = 10 | 2000/1000<br>n = 10 | control<br>n = 10 | 20/10<br>n = 10 | 200/100<br>n = 10 | 2000/1000<br>n = 10 |
| Glu (mmol/L) | 7.39 ± 0.73 | 7.10 ± 0.85 | 6.93 ± 0.61 | 7.10 ± 0.61 | 6.00 ± 1.14 | 6.04 ± 0.83 | 5.88 ± 0.46 | 6.22 ± 0.86 |
| BUN (mmol/L) | 5.72 ± 1.01 | 6.00 ± 0.51 | 5.78 ± 0.95 | 5.91 ± 0.48 | 7.25 ± 0.91 | 6.86 ± 1.13 | 7.67 ± 0.86 | 7.76 ± 1.08 |
| Creat (μmol/L) | 44 ± 4 | **\*53 ± 7** | **\*51 ± 4** | **\*53 ± 5** | 54 ± 9 | 54 ± 7 | 58 ± 6 | 50 ± 6 |
| AST (U/L) | 87 ± 12 | 105 ± 33 | 97 ± 15 | 88 ± 8 | 107 ± 23 | 103 ± 19 | 99 ± 15 | 84 ± 17 |
| ALT (U/L) | 30 ± 7 | 30 ± 7 | 31 ± 7 | 25 ± 5 | 26 ± 5 | 27 ± 4 | 25 ± 5 | 31 ± 9 |
| GGT (U/L) | 4 ± 0 | 4 ± 0 | 4 ± 0 | 4 ± 1 | 4 ± 0 | 4 ± 0 | 4 ± 1 | 4 ± 1 |
| Alp (U/L) | 80 ± 12 | 85 ± 8 | 85 ± 12 | 84 ± 11 | 38 ± 9 | 36 ± 7 | 37 ± 11 | 47 ± 12 |
| LDH (U/L) | 170 ± 45 | 181 ± 44 | 163 ± 39 | 208 ± 44 | 225 ± 47 | 193 ± 40 | 212 ± 33 | **\*\*171 ± 48** |
| T.Bil (μmol/L) | 3.02 ± 0.51 | 2.90 ± 0.68 | 2.95 ± 0.42 | **\*\*1.41 ± 0.30** | 3.21 ± 0.74 | 3.03 ± 0.51 | **\*\*2.39 ± 0.87** | **\*\*1.69 ± 0.55** |
| T.Chol (mmol/L) | 1.79 ± 0.17 | 1.73 ± 0.26 | 1.74 ± 0.22 | 1.72 ± 0.14 | 1.96 ± 0.27 | 2.05 ± 0.33 | 2.14 ± 0.31 | 2.10 ± 0.36 |
| AHDL (mmol/L) | 1.46 ± 0.14 | 1.44 ± 0.23 | 1.43 ± 0.16 | 1.39 ± 0.14 | 1.61 ± 0.21 | 1.72 ± 0.26 | 1.75 ± 0.23 | 1.66 ± 0.28 |
| LDL Cholesterol (mmol/L) | 0.21 ± 0.07 | **\*\*0.13 ± 0.05** | 0.17 ± 0.07 | 0.22 ± 0.04 | 0.28 ± 0.06 | 0.25 ± 0.09 | 0.30 ± 0.08 | 0.34 ± 0.09 |
| Trig(mmol/L) | 0.63 ± 0.29 | 0.80 ± 0.46 | 0.68 ± 0.10 | 0.60 ± 0.20 | 0.35 ± 0.09 | 0.41 ± 0.14 | 0.45 ± 0.15 | 0.48 ± 0.16 |
| T.Pro(g/L) | 68.6 ± 2.0 | **\*71.4 ± 3.0** | **\*73.1 ± 2.0** | 71.0 ± 2.4 | 78.1 ± 3.3 | 79.7 ± 2.9 | **\*81.7 ± 2.2** | 79.7 ± 2.2 |
| ALB(g/L) | 30.3 ± 0.6 | 30.1 ± 0.7 | **\*34.2 ± 1.6** | **\*36.6 ± 0.6** | 37.7 ± 1.3 | 37.4 ± 1.1 | **\*44.1 ± 1.1** | **\*44.6 ± 1.5** |
| GLOB(g/L) | 38.4 ± 1.6 | **\*41.3 ± 2.9** | 39.0 ± 1.9 | **\*\*34.5 ± 2.2** | 40.4 ± 2.3 | 42.4 ± 2.1 | **\*\*37.7 ± 1.6** | **\*\*35.2 ± 1.3** |
| A/G(ratio) | 0.79 ± 0.03 | **\*\*0.73 ± 0.06** | **\*0.88 ± 0.07** | **\*1.06 ± 0.06** | 0.94 ± 0.04 | **\*\*0.88 ± 0.03** | **\*1.17 ± 0.04** | **\*1.27 ± 0.06** |
| Pi(mmol/L) | 2.12 ± 0.32 | 2.03 ± 0.32 | 1.89 ± 0.34 | 2.04 ± 0.57 | 1.75 ± 0.33 | 1.75 ± 0.31 | 2.34 ± 2.14 | 1.86 ± 0.65 |
| Ca(mmol/L) | 2.85 ± 0.03 | 2.80 ± 0.08 | **\*\*2.69 ± 0.15** | **\*\*2.70 ± 0.07** | 2.88 ± 0.07 | 2.89 ± 0.07 | 2.75 ± 0.15 | 2.89 ± 0.09 |
| Na(mEq/L) | 143.3 ± 0.6 | **\*144.3 ± 0.8** | **\*145.8 ± 1.1** | 143.6 ± 0.9 | 142.4 ± 1.1 | 143.5 ± 1.3 | **\*145.4 ± 1.4** | 143.2 ± 0.8 |
| K(mEq/L) | 3.78 ± 0.16 | 3.95 ± 0.24 | 3.65 ± 0.24 | **\*4.28 ± 0.25** | 3.48 ± 0.48 | 3.55 ± 0.30 | 3.56 ± 0.24 | **\*4.01 ± 0.36** |
| Cl(mEq/L) | 98.4 ± 0.8 | 98.7 ± 0.9 | 98.8 ± 1.2 | 97.8 ± 0.8 | 96.6 ± 1.5 | 96.7 ± 1.5 | 97.8 ± 1.3 | 96.8 ± 1.1 |

Values are expressed as mean ± standard deviation. mg/kg/day, milligrams per kilogram body weight per day; L, liter; n, number of animals, U, ;mEq, ; g, gram; \*/\*\* Significantly higher/lower than the vehicle control group at p < 0.05

The creatinine increase at all doses in males was minimal and was not dose dependent (Table 4). A dose dependent decrease in total bilirubin for females was noted, however, in the absence of significant changes in red blood cell parameters, the decreased total bilirubin values were considered to be toxicologically insignificant. There were no corresponding microscopic findings at the high dose to indicate this as test item-related. All the other intergroup differences were also considered incidental as the changes were not dose-related. Further, the individual animal values were within the range of normal variation.

Thyroid hormone profile (T3, T4 and TSH) was not affected by the test item administration. In males, an increase in TSH level was noted at the mid dose (Table 5). As there were no corresponding changes at the high dose, this finding was not considered as test item-related. The urine parameters were not affected by test item administration.

**TABLE 5.** Summary of Significant Results in the Thyroid Hormone Profile.

|  | Dose Group (mg/kg/day) |  |  |  |
| --- | --- | --- | --- | --- |
|  | Males |  |  |  |
|  | control<br>n = 10 | 20/10<br>n = 10 | 200/100<br>n = 10 | 2000/1000<br>n = 10 |
| T3 (ng/mL) | 0.47 ± 0.14 (--) | 0.54 ± 0.15 (--) | 0.6 ± 0.31 (--) | 0.52 ± 0.12 (--) |
| T4 (ng/mL) | 24.55 ± 9.27 (--) | 28.01 ± 12.5 (--) | 29.1 ± 10.84 (--) | 30.58 ± 14.08 (--) |
| TSH (ng/mL) | 0.89 ± 0.44 (--) | 0.96 ± 0.44 (--) | **\*1.95 ± 0.74 (119)** | 1.21 ± 0.51 (--) |

Values are expressed as mean ± standard deviation. mg/kg/day, milligrams per kilogram body weight per day; ng/mL, nanograms per millilitre; \* Significantly higher than the vehicle control group at p < 0.05, percent change given in brackets; (--) percent change not presented as difference is not significant.

The terminal fasting body weights were not affected by test item administration, and there were no test item-related changes in the organ weights and their ratios. A minimal increase in liver weight (relative to body weight-9 % and relative to brain weight-14 %) was observed in males at high dose. However, these rats did not show any changes in the absolute weight of these organs and there were no associated biochemical or microscopic changes. Thus this finding was not considered as test item-related.

All the other organ weight changes were also considered incidental as the differences were very minimal and not dose related.

There were no test item-related gross lesions or microscopic changes. The incidences of different findings at the high dose were similar to concurrent vehicle control group and were the common spontaneous changes for this age group rats.

## Conclusion

The results of the study indicate that the oral gavage administration of the lyophilized *B. subtilis* ZB183 spores at the doses of 10^9^, 10^10^, and 10^11^ CFU/kg body weight/day to Wistar rats for 90 consecutive days had no effects on clinical signs, mortality, ophthalmological examinations, functional observational battery, body weights, body weight gains and food consumption in both sexes. There were no test item-related changes observed in haematology, coagulation, urinalysis, thyroid hormonal analysis (T3, T4 and TSH), terminal fasting body weights, organ weights, gross pathology and histopathology. An isolated incidence of minimal increase in the plasma albumin level at 10^10^ and 10^11^ CFU/kg/day doses without an increase in total protein was considered non-adverse effect. An incidence of decrease in total bilirubin values for females at 10^10^ and 10^11^ CFU/kg/day doses was considered to be toxicologically insignificant in the absence of significant changes in red blood cell parameters.

Thus, the lyophilized *B. subtilis* ZB183 spores did not show any toxicological effects when administered orally for 90 consecutive days in Wistar rats at all the tested doses. Hence, the “No Observed Adverse Effect Level (NOAEL)” is defined at 10^11^ CFU/kg body weight/day for lyophilized *B. subtilis* ZB183 spores under the test conditions employed.

## Competing Interests

ZBiotics Company intends to use *B. subtilis* ZB183 as an ingredient in products for sale and stands to benefit financially if *B. subtilis* ZB183 is determined to be safe. ZBiotics Company contracted Eurofins Advinus to perform the 90-day toxicity study discussed in this paper. ZBiotics Company did not participate in any of the experimentation, evaluation, or analysis, but the authors listed from ZBiotics Company did assist in assembling this paper. However, to ensure an unbiased interpretation of the data, Eurofins Advinus wrote all of the results and discussion sections with only compilation and minimal linguistic editing by ZBiotics, and final approval of the accuracy was given by Eurofins Advinus. Otherwise, Eurofins Advinus has no financial or other interest in ZBiotics Company or *B. subtilis* ZB183 beyond the contracted work described in this paper.

## Funding Statement

This work was paid for by ZBiotics Company, which did participate in the compilation, editing, and submission of this manuscript.

## Acknowledgements

We would like to thank Dr. Claire Kruger and ChromaDex Spherix Consulting (11821 Parklawn Drive, Suite 310 Rockville, MD 20852) for their help and input in designing the study. In addition, we would like to thank Dr. John Endres at AIBMR Life Sciences, Inc. (2800 E. Madison St., Suite 202 - Seattle, WA 98112) for reviewing the manuscript and providing feedback.

